## Supplemental Figures TableS1, FiguresS1-S13 for "Conformation Selection by ATP-competitive Inhibitors and Allosteric Communication in ERK2"

**SUPPLEMENTAL TABLE**

Table S1. X-ray data collection and refinement parameters

**SUPPLEMENTAL FIGURES**

Figure S1. Structural features of ERK2.
Figure S2. 2D-HMQC NMR spectra of apoenzyme and inhibitor-bound ERK2.
Figure S3. Titration of inhibitor binding to 2P-ERK2.
Figure S4. Proteolytic peptides analyzed by HDX-MS.
Figure S5. Structural map of regions showing HDX responses to inhibitor binding.
Figure S6. HDX time courses of regions with comparable responses to different inhibitors.
Figure S7. HDX time courses of differential responses to inhibitor binding.
Figure S8. Differential HDX responses to new inhibitors surveyed in this study.
Figure S9. 2D-HMQC NMR spectra of inhibitor-bound 2P-ERK2.
Figure S10. Chemical shift changes induced by inhibitor binding to 2P-ERK2.
Figure S11. HDX time courses responsive to ATG017 binding.
Figure S12. Interactions of L16 and helix αL16 with N-lobe elements.
Figure S13. Crystal contacts with the activation loop in X-ray structures of ERK2

**SUPPLEMENTAL DATASETS**

Dataset S1. Deuterium uptake time courses for all HDX-MS experiments (Excel format)
Dataset S2. NMR chemical shifts and perturbations by selected inhibitors (Excel format)

**SUPPLEMENTAL TABLE**

**Table S1. X-ray data collection and refinement statistics**

| | **2P-ERK2:Inhibitor #8** | **2P-ERK2:Inhibitor #16** |
|---|---|---|
| **PDB code** | 8U8K | 8U8J |
| **Data Collection:** | | |
| Space group | $P2_12_12_1$ | $P2_12_12_1$ |
| Unit cell parameters (Å,°) | a = 41.98, b = 77.28,<br>c = 151.54, β = 90.0 | a = 42.10, b = 76.66,<br>c = 151.97, β = 90.0 |
| Resolution (Å) | 21.30 – 2.10<br>2.17 – 2.10 | 19.56 – 2.10<br>2.21 – 2.10 |
| Unique reflections (last shell) | 29,455 | 25,560 |
| Data Completeness (%) | 99.5 (21.30 – 2.10) | 86.5 (19.56 – 2.10) |
| Data Redundancy | 6.50 | 5.66 |
| $R_{sym}$ | 0.15 | 0.15 |
| $< I/\sigma(I) >$ | 4.60 (at 2.09 Å) | 3.66 (at 2.09 Å) |
| Refinement program | PHENIX 1.20.1_4487 | PHENIX 1.20.1_4487 |
| $R_{work}$, $R_{free}$ | 0.174 , 0.213 | 0.177 , 0.227 |
| $R_{free}$ test set | 1497 reflections (5.08%) | 1299 reflections (5.08%) |
| Wilson B-factor (Å$^2$) | 25.9 | 27.4 |
| Anisotropy | 0.116 | 0.125 |
| $F_o$, $F_c$ correlation | 0.95 | 0.95 |
| Total residues | 349 | 352 |
| Total atoms, non-hydrogen | 3,138 | 3,131 |
| Protein atoms, non-hydrogen | 2,858 | 2,873 |
| Ligand heteroatoms | 30 | 27 |
| Water molecules | 250 | 231 |
| Bond length RMSZ<br># \|Z\| > 5 | 0.37<br>0 / 2898 | 0.38<br>0 / 2913 |
| Bond angles RMSZ<br># \|Z\| > 5 | 0.56<br>0 / 3924 | 0.57<br>1 / 3944 |
| Average B, all atoms (Å$^2$) | 27.0 | 27.0 |

### Figure S1

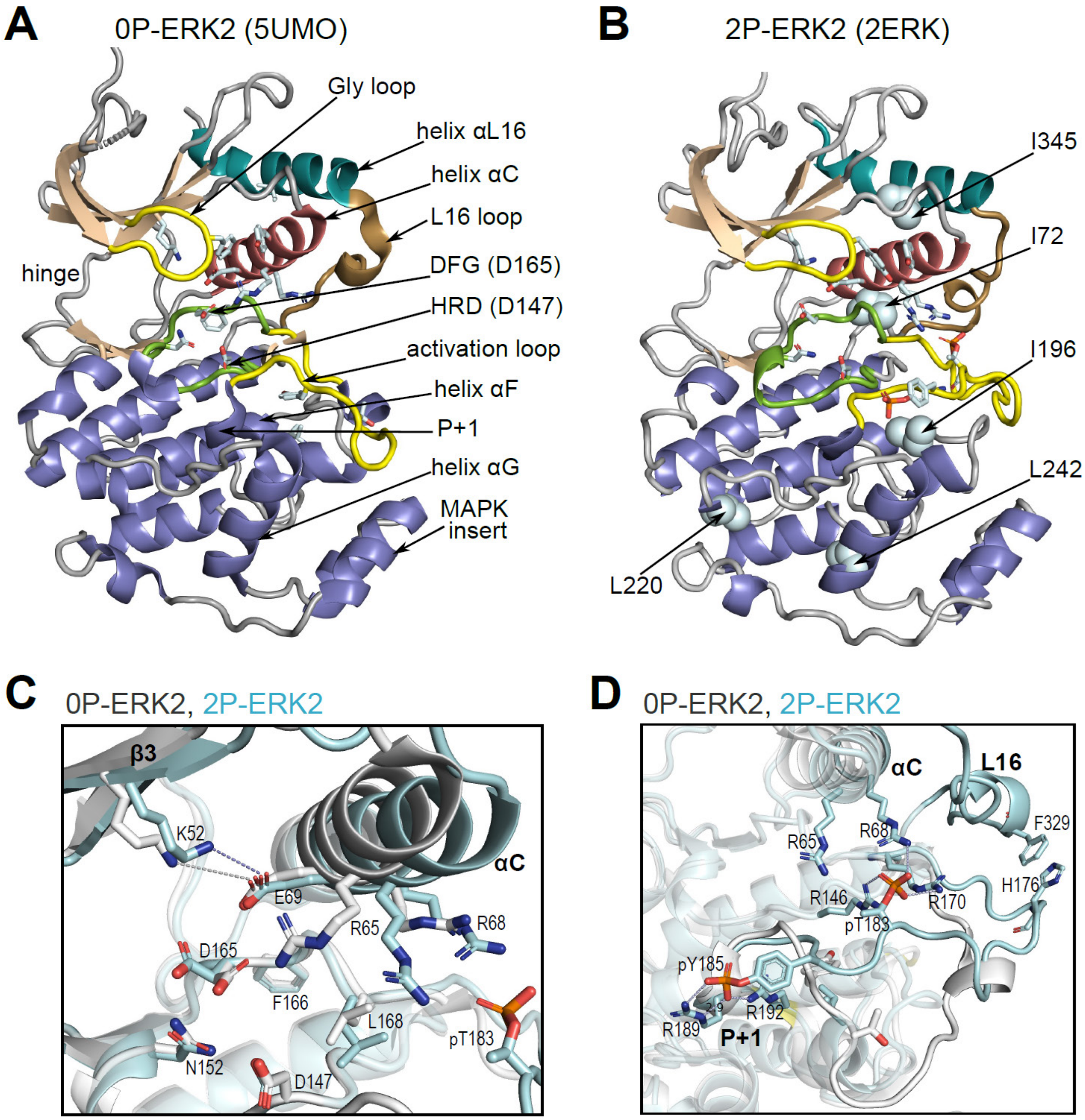


**Figure S1. Structural features of ERK2. (A,B)** X-ray structures of **(A)** 0P-ERK2 (PDBID:5UMO) and **(B)** 2P-ERK2 (PDBID:2ERK) apoenzymes. Panel **A** labels conserved motifs in ERK2 common to protein kinases. Panel **B** labels residues that illustrate L⇌R exchange (I72, L220, L242; **Fig. 1** and **Fig. 6**), and key chemical shift perturbations (I196, I345; **Fig. 7**). **(C)** Structural superposition of 0P-ERK2 (white) and 2P-ERK2 (pale blue), illustrating overlapping positions of active site residues and an outward shift of helix αC upon dual phosphorylation. Root-mean-square deviations between 0P- and 2P-ERK2 for catalytic site residues were K52-0.69 Å, E69-0.16 Å, D147-0.055 Å, D165-0.88 Å. **(D)** Superposition of 0P-ERK2 and 2P-ERK2, illustrating conformational differences in the activation loop. In 2P-ERK2, pT183 and pY185 form ion pairs with multiple Arg residues, while the L16 loop folds into a 3/10 helix with side chain interactions to the activation loop. Structures were superpositioned by aligning Cα atoms within the C-terminal domain (residues 109-141, 205-245, 272-310).

### Figure S2

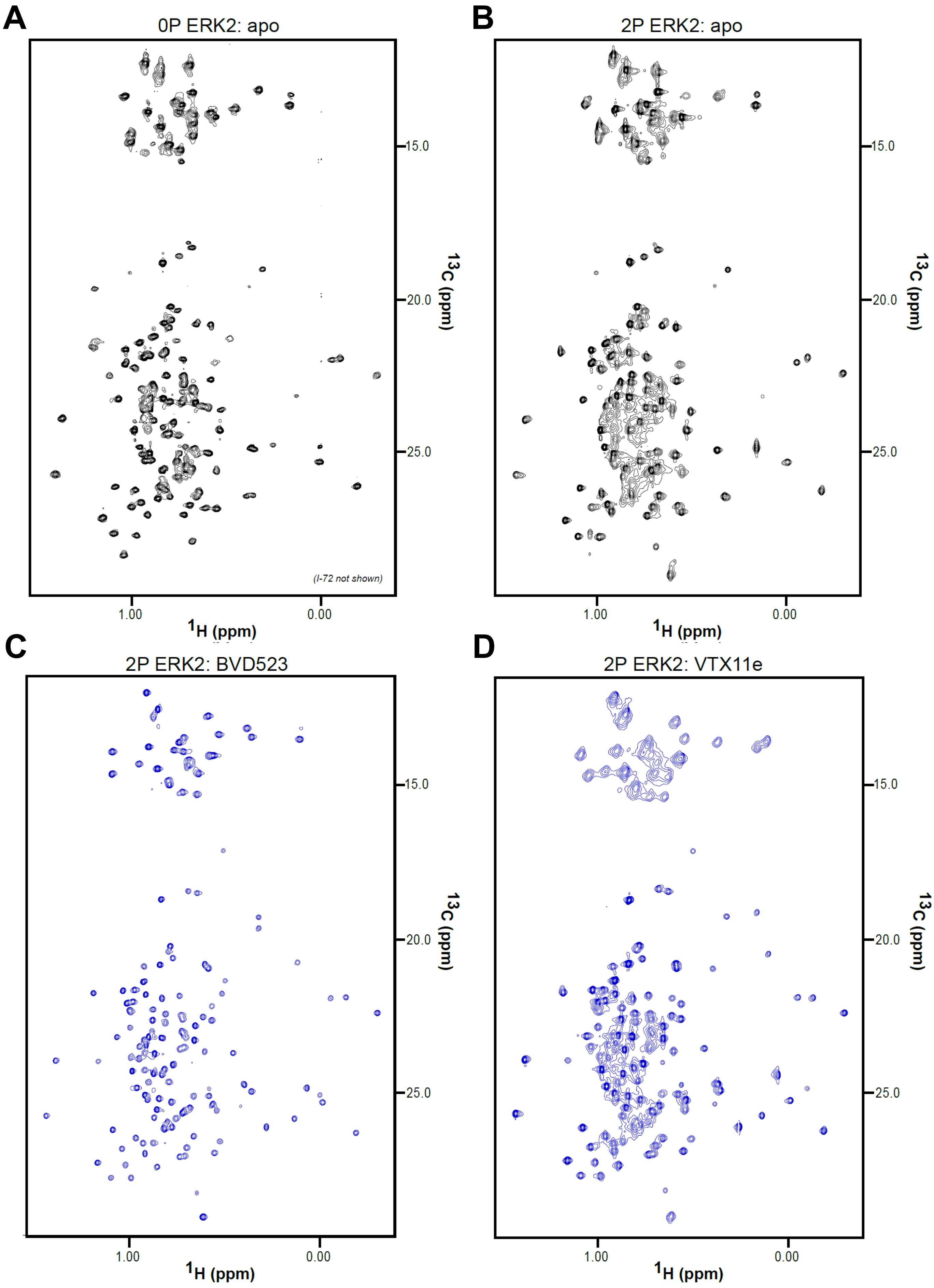

A
0P ERK2: apo
(I-72 not shown)
B
2P ERK2: apo
C
2P ERK2: BVD523
D
2P ERK2: VTX11e
15.0
20.0
25.0
13C (ppm)
1.00
0.00
1H (ppm)

### Figure S2

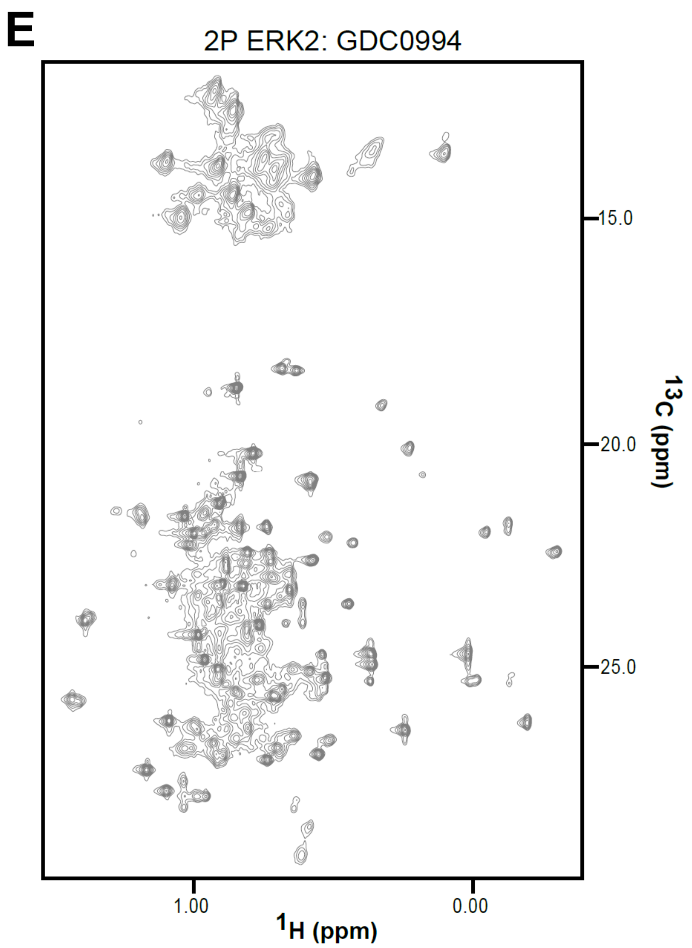


**Figure S2. 2D-HMQC NMR spectra of apoenzyme and inhibitor-bound ERK2.** 2D [$^{13}$C,$^{1}$H] HMQC spectra for **(A)** apo 0P-ERK2, **(B)** apo 2P-ERK2, **(C)** 2P-ERK2:BVD523, **(D)** 2P-ERK2:VTX11e, and **(E)** 2P-ERK2:GDC0994. Each complex was formed from 150 μM ERK2 and 180 μM inhibitor, yielding binding saturation at concentration ratios [2P-ERK2]:[inhibitor] = 1.0:1.2.

#### Figure S3A

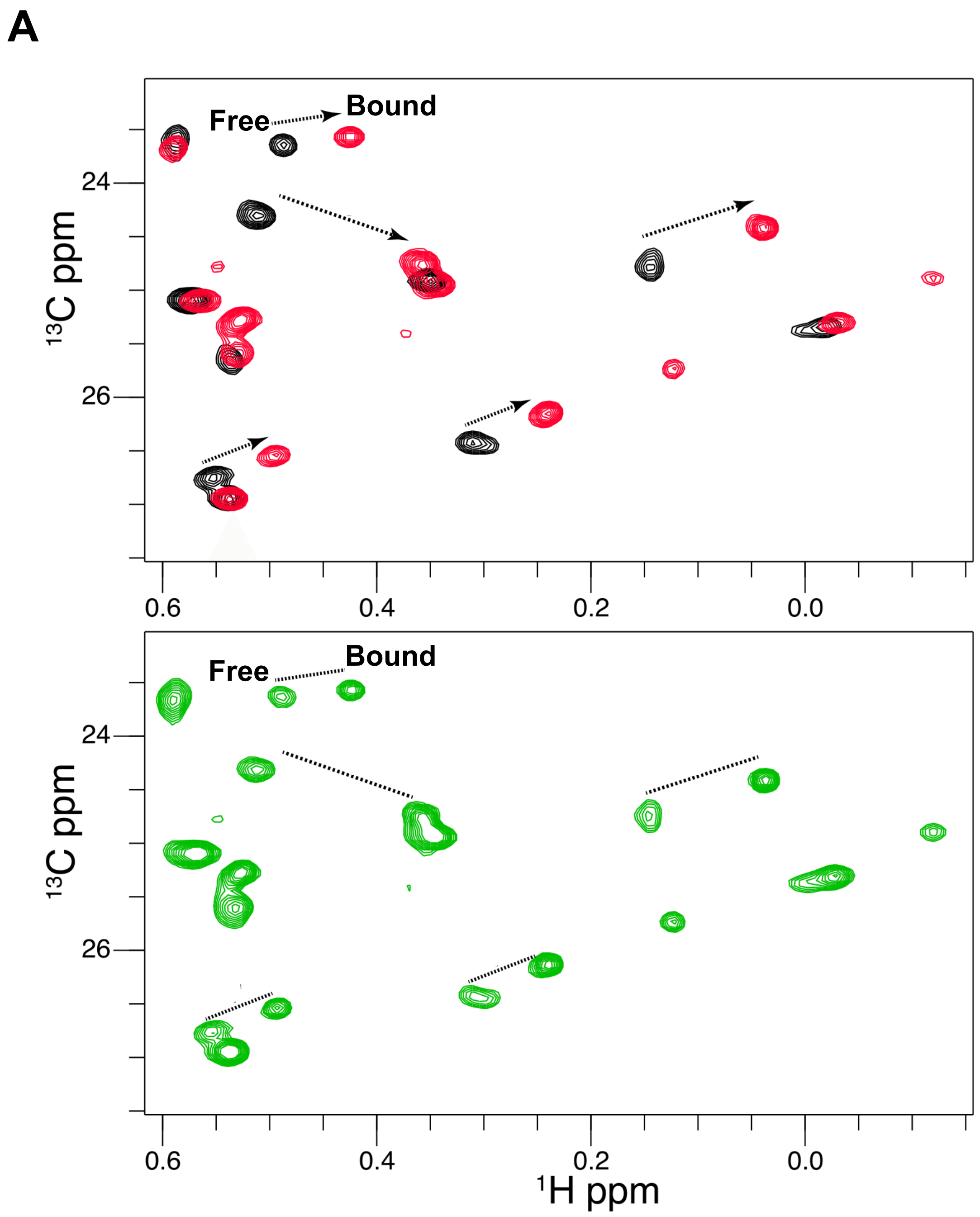
A
Free
Bound
24
26
13C ppm
0.6
0.4
0.2
0.0
Free
Bound
24
26
13C ppm
0.6
0.4
0.2
0.0
1H ppm

#### Figure S3B

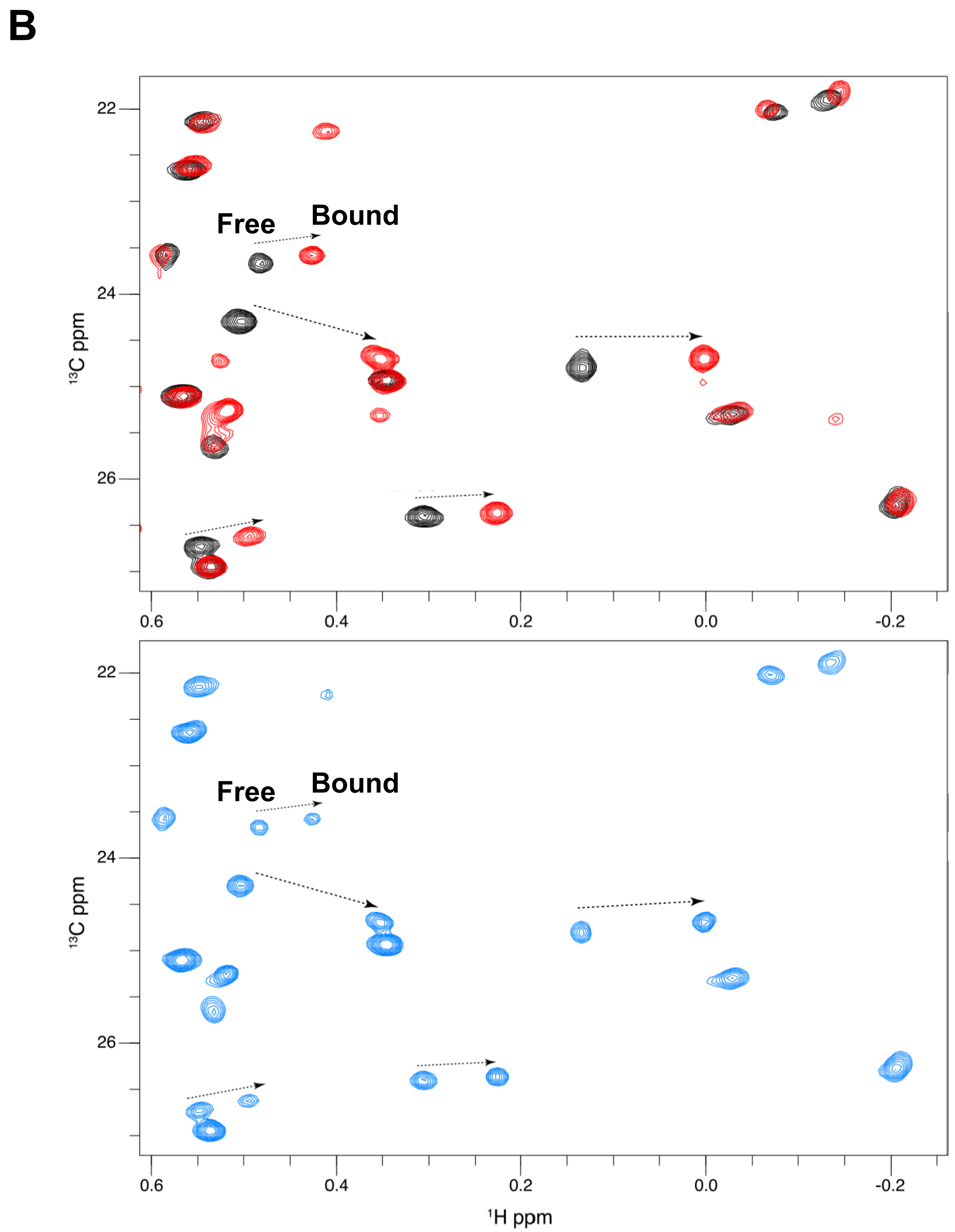


**Figure S3. Titration of inhibitor binding to 2P-ERK2.** 2D [$^{13}$C,$^{1}$H] HMQC spectra reveal residues in slow exchange that report saturation binding with increasing inhibitor concentration. **(A)** NMR spectra of VTX11e complexed with 2P-ERK2. Black: 2P-ERK2 apoenzyme; Green: 50% bound, [2P-ERK2]:[VTX11e] = 1.0:0.5; Red: 100% bound, [2P-ERK2]:[VTX11e] = 1.0:1.3. **(B)** NMR spectra of GDC0994 complexed with 2P-ERK2. Black: 2P-ERK2 apoenzyme; Blue: 50% bound, [2P-ERK2]:[GDC0994] = 1.0:0.5; Red: 100% bound, [2P-ERK2]:[GDC0994] = 1.0:1.3. Arrows show the shifts between free and bound forms of methyl peaks. The results illustrate that binding saturation is confirmed when peaks reflecting unbound ERK2 are undetectable.

### Figure S4

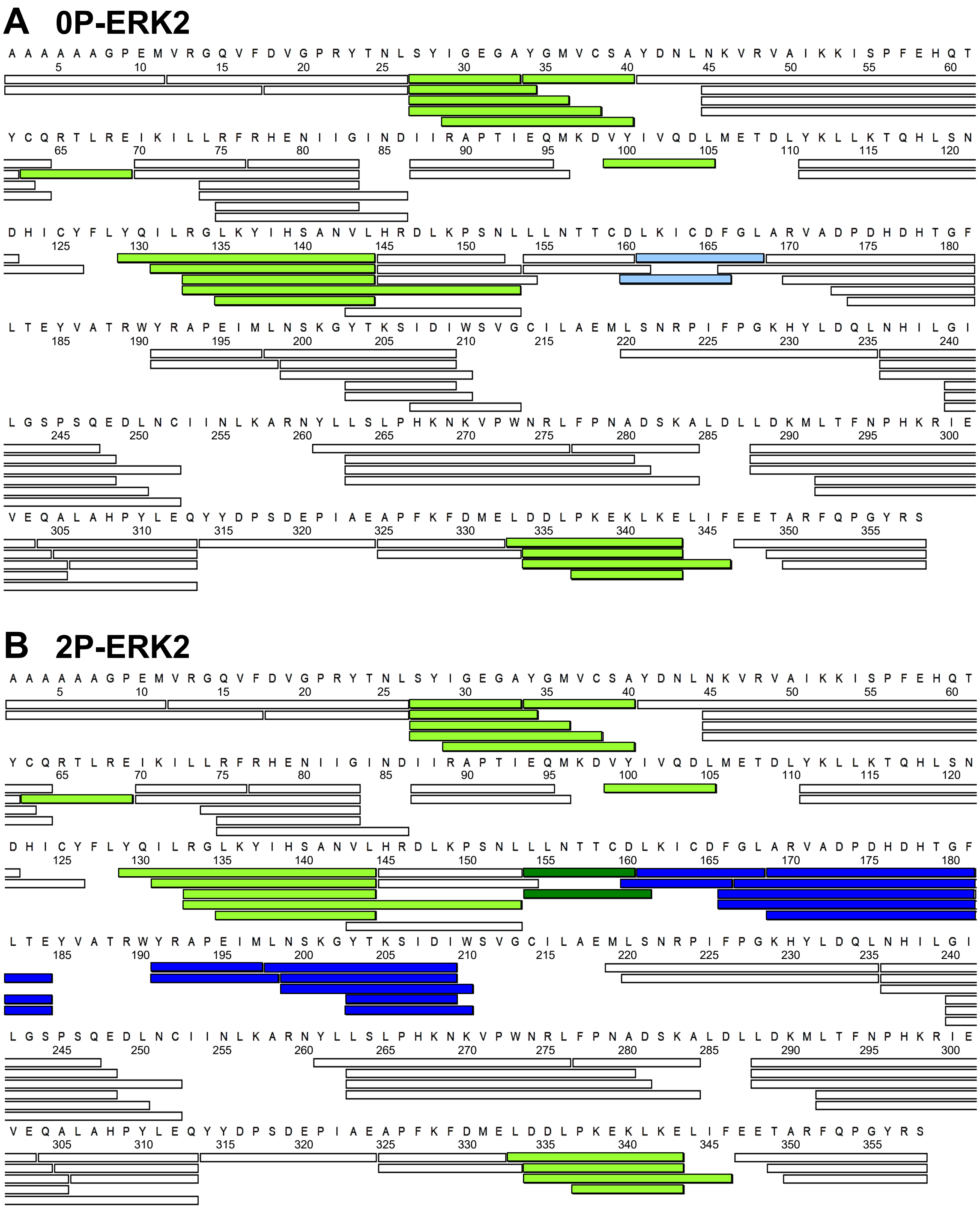


**Figure S4. Proteolytic peptides analyzed by HDX-MS.** Peptides produced by pepsin digestion yielded 90% and 91% coverage of exchangeable amides in **(A)** 0P-ERK2 and **(B)** 2P-ERK2, respectively. Colors indicate regions where ligand binding alters HDX uptake. Light green indicates peptides where binding of VTX11e, BVD523, and GDC0994 induce a similar degree of HDX protection (i.e., decreased deuterium uptake) in both 0P-ERK2 and 2P-ERK2. Deep green indicates peptides where all inhibitors induce a similar degree of HDX protection, but only in 2P-ERK2. Deep blue indicates peptides where VTX11e and BVD523 induce greater HDX protection compared to GDC0994 in 2P-ERK2. Light blue indicates peptides that show HDX protection by all inhibitors around the DFG motif in 0P-ERK2, but to a lower amount compared to BVD523 or VTX11e in 2P-ERK2.

### Figure S5

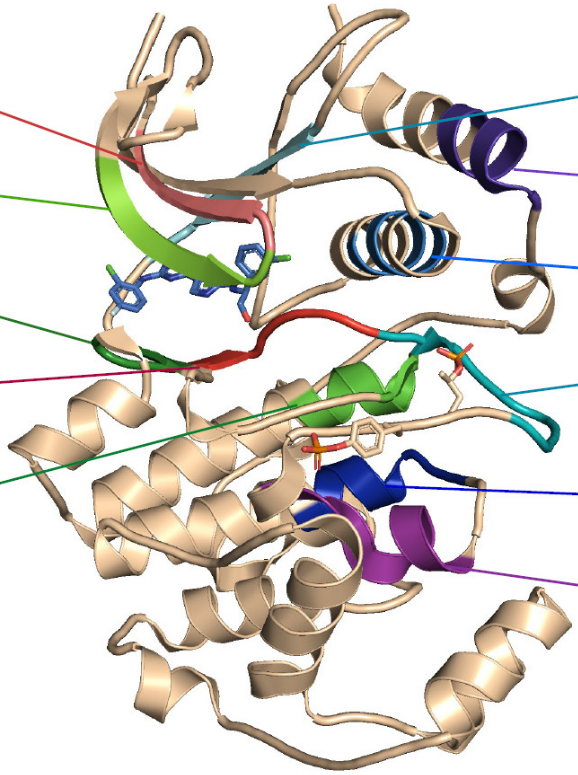


**Figure S5. Structural map of regions showing HDX responses to inhibitor binding.** Structure of 2P-ERK2 (PDBID:6OPK), indicating the locations of peptides where binding of VTX11e, BVD523 or GDC0994 alter HDX behavior, as highlighted in **Suppl. Fig. S4**.

### Figure S6

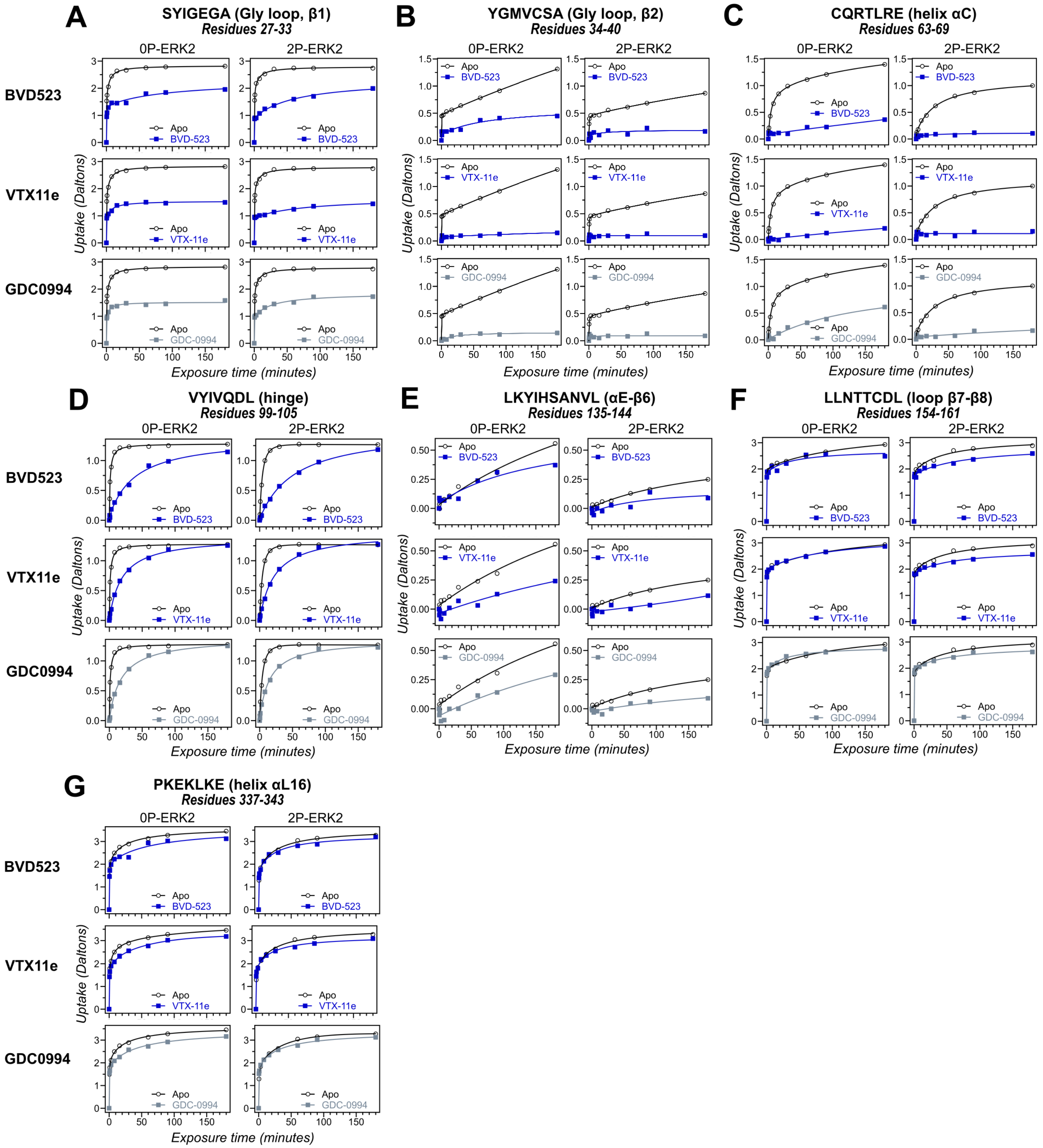


**Figure S6. HDX time courses of regions with comparable responses to different inhibitors.** HDX time courses for key peptide segments in ERK2, including **(A,B)** the Gly loop (peptide 29-33, SYIGEGA; peptide 34-40, YGMVCSA), **(C)** helix αC (peptide 63-69, CQRTLRE), **(D)** the hinge (peptide 99-105, VYIVQDL), **(E)** helix αE-β6 (peptide 135-144, LKYIHSANVL), **(F)** loop β7-β8 (peptide 154-161, LLNTTCDL), and **(G)** helix αL16 (peptide 337-343, PKEKLKE). Open symbols show deuterium uptake in 0P- or 2P-ERK2 apoenzymes. Closed symbols show deuterium uptake in 0P- or 2P-ERK2 complexed with BVD523, VTX11e (blue) or GDC0994 (grey). In these regions, deuterium uptake decreases by a similar degree upon binding each of the three inhibitors.

#### Figure S7

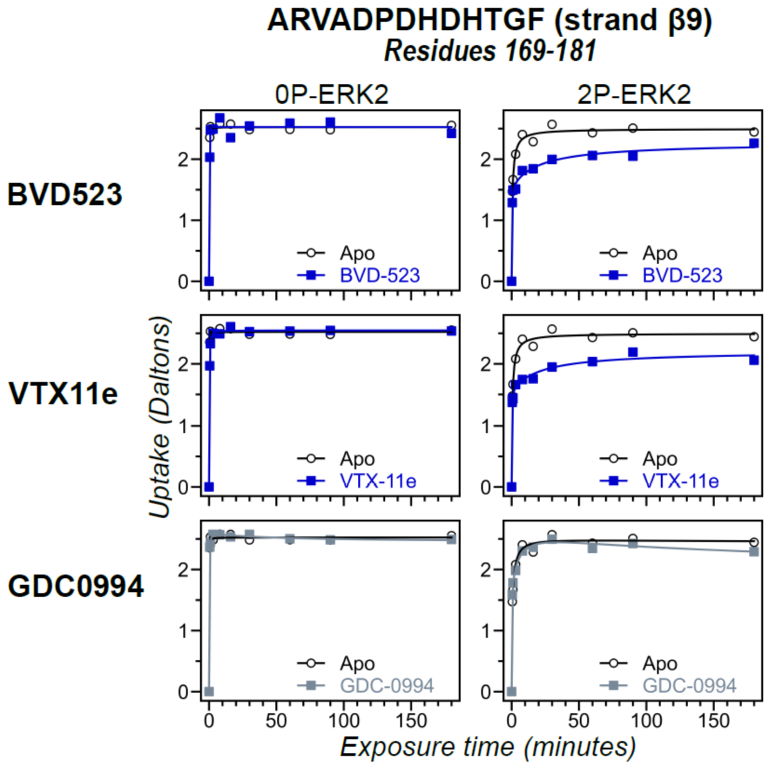


**Figure S7. HDX time courses with differential responses to inhibitor binding.** HDX time courses for strand β9 (peptide 169-181, ARVADPDHDHTGF) located adjacent to the DFG motif. Open symbols show deuterium uptake in 0P- or 2P-ERK2 apoenzymes. Closed symbols show deuterium uptake in 0P- or 2P-ERK2 complexed with BVD523, VTX11e or GDC0994. Deuterium uptake in these regions decreases by a larger degree upon binding VTX11e or BVD523 (blue) compared to GDC0994 (grey), as also seen in the DFG motif, P+1 and helix αF (**Fig. 2**).

### Figure S8A

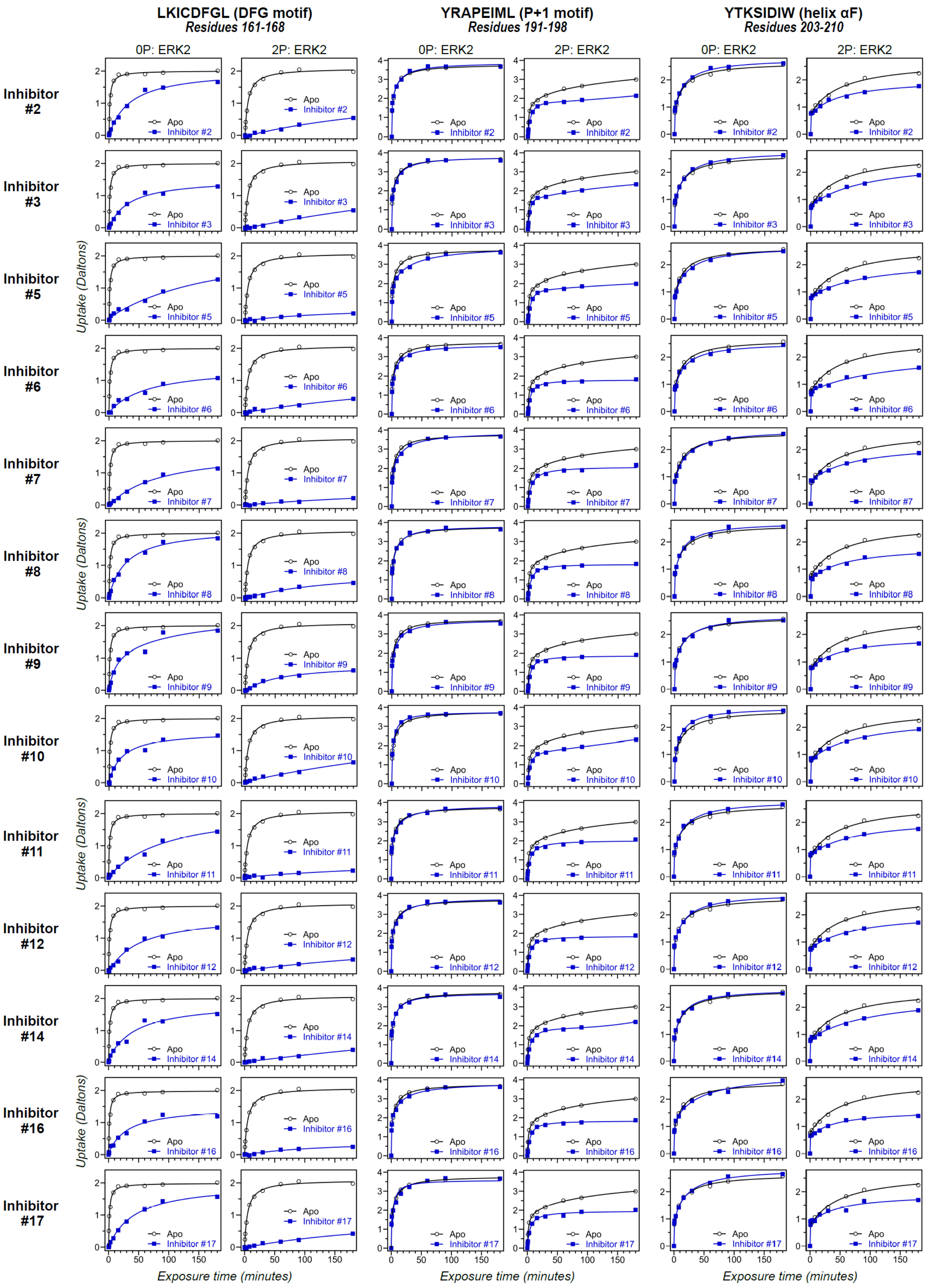

LKICDFGL (DFG motif)
Residues 161-168
YRAPEIML (P+1 motif)
Residues 191-198
YTKSIDIW (helix αF)
Residues 203-210
0P: ERK2
2P: ERK2
Inhibitor #2
Inhibitor #3
Inhibitor #5
Inhibitor #6
Inhibitor #7
Inhibitor #8
Inhibitor #9
Inhibitor #10
Inhibitor #11
Inhibitor #12
Inhibitor #14
Inhibitor #16
Inhibitor #17
Apo
Uptake (Daltons)
Exposure time (minutes)

### Figure S8B

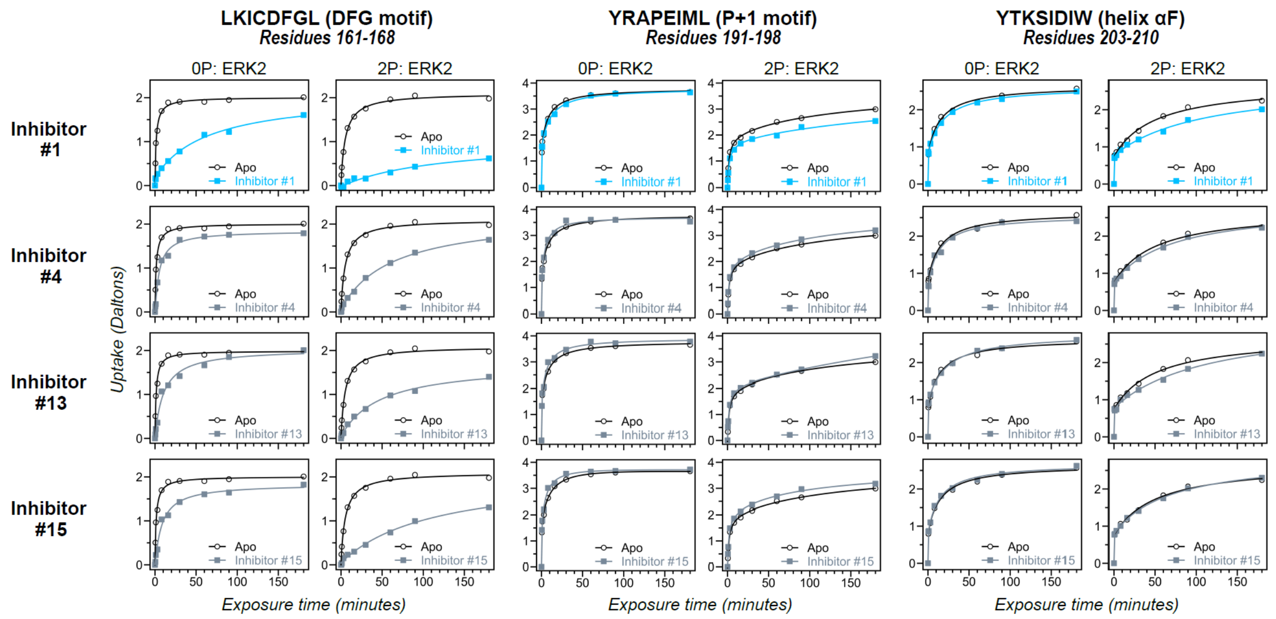


**Figure S8. Differential HDX responses to new inhibitors surveyed in this study.** Effects of binding the full set of inhibitors #1-#17 on deuterium uptake into the DFG motif (peptide 161-168: LKICDFGL), P+1 segment (peptide 191-198, YRAPEIML), and helix αF (peptide 203-210: YTKSIDIW), expanding the selected subset in **Fig. 4**. Open symbols show deuterium uptake in 0P- or 2P-ERK2 apoenzymes. Closed symbols show deuterium uptake in 0P- or 2P-ERK2 complexed with inhibitors. **(A)** Thirteen inhibitors (#2, #3, #5-12, #14, #16 and #17; blue) show HDX uptake patterns for 2P-ERK2 that are similar to VTX11e and BVD523, suggesting conformation selection for the R-state. **(B)** Three inhibitors (#4, #13, #15; grey) show HDX patterns similar to GDC0994, consistent with R⇌L exchange. One inhibitor (#1; cyan) shows significant HDX protection in the DFG motif, but only marginal protection in the P+1 segment or helix αF, suggesting behavior intermediate between VTX11e/BVD523 and GDC0994.

#### Figure S9A-D

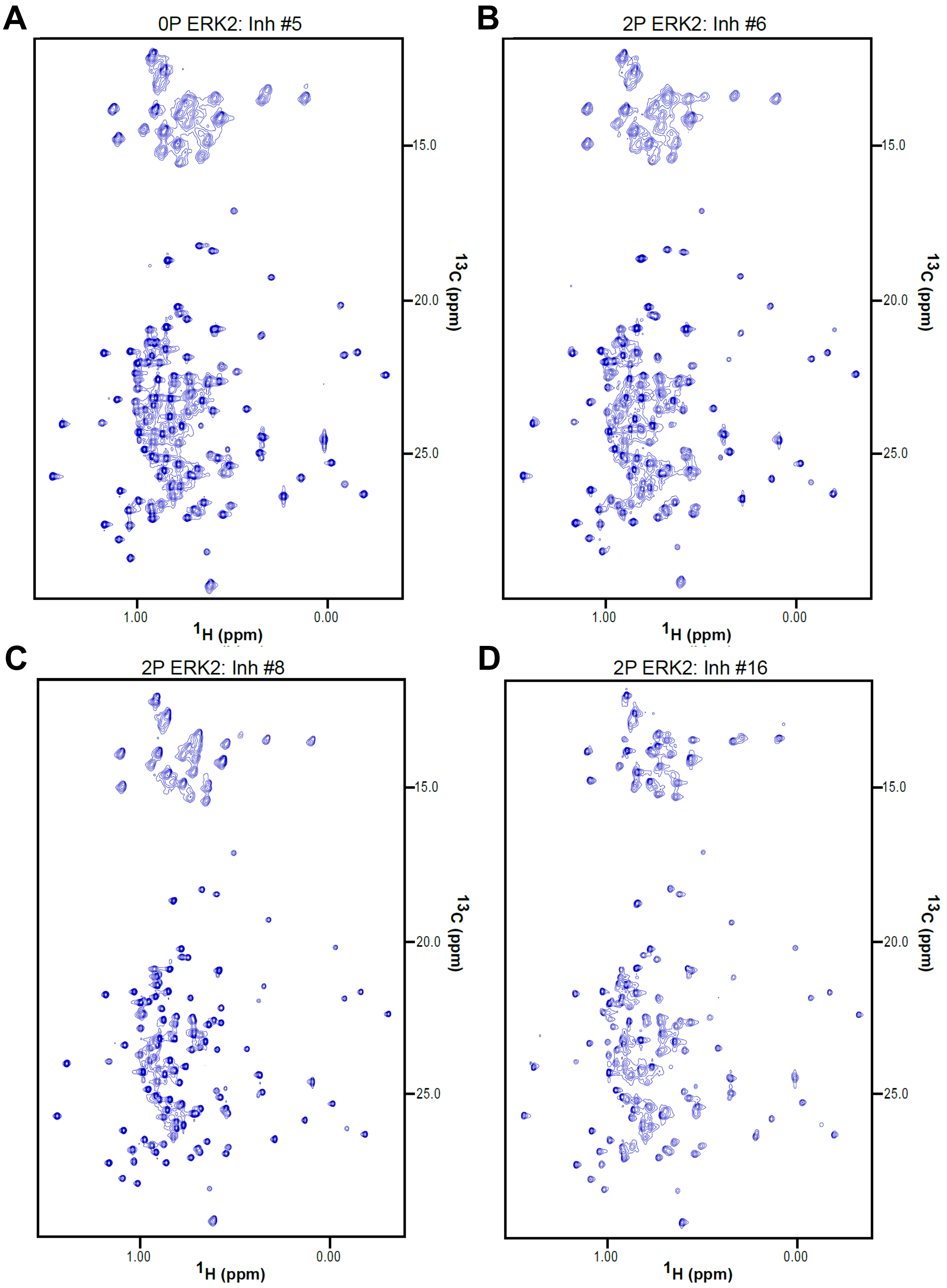

A
0P ERK2: Inh #5
B
2P ERK2: Inh #6
C
2P ERK2: Inh #8
D
2P ERK2: Inh #16
15.0
20.0
25.0
13C (ppm)
1.00
0.00
1H (ppm)

### Figure S9E-G

E

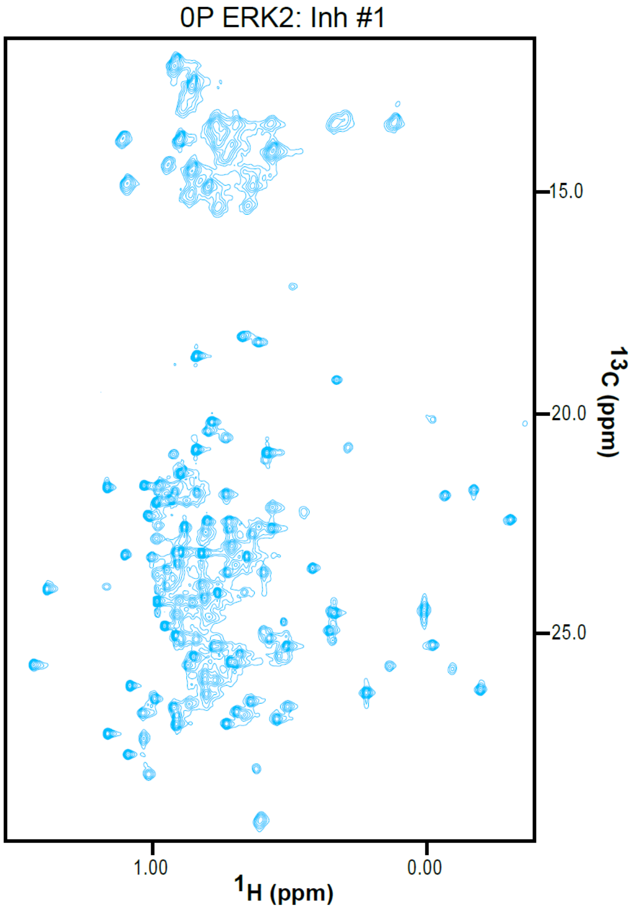


F

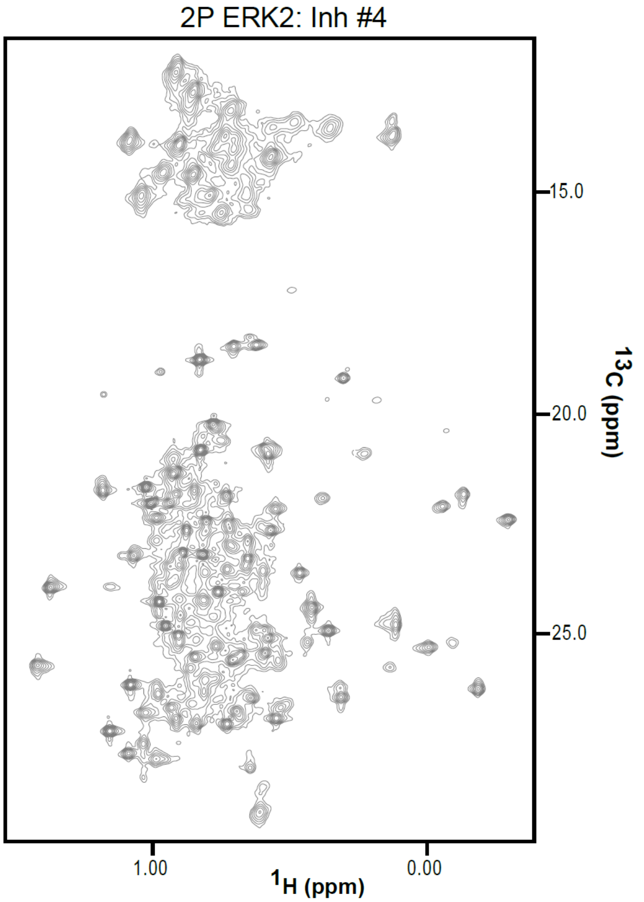


G

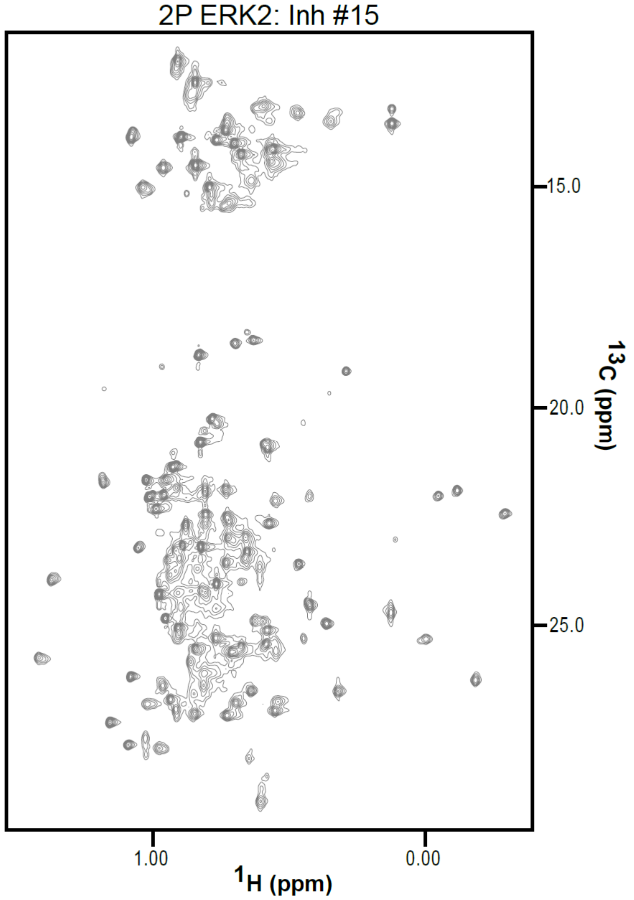

### Figure S9

**Figure S9. 2D-HMQC NMR spectra of inhibitor-bound 2P-ERK2.** 2D [$^{13}C$,$^{1}H$] HMQC spectra for **(A)** 2P-ERK2:Inh#5, **(B)** 2P-ERK2:Inh#6, **(C)** 2P-ERK2:Inh#8, **(D)** 2P-ERK2:Inh#16, **(E)** 2P-ERK2:Inh#1, **(F)** 2P-ERK2:Inh#4, and **(G)** 2P-ERK2:Inh#15. Each complex was formed from 150 μM ERK2 and 180 μM inhibitor.

### Figure S10

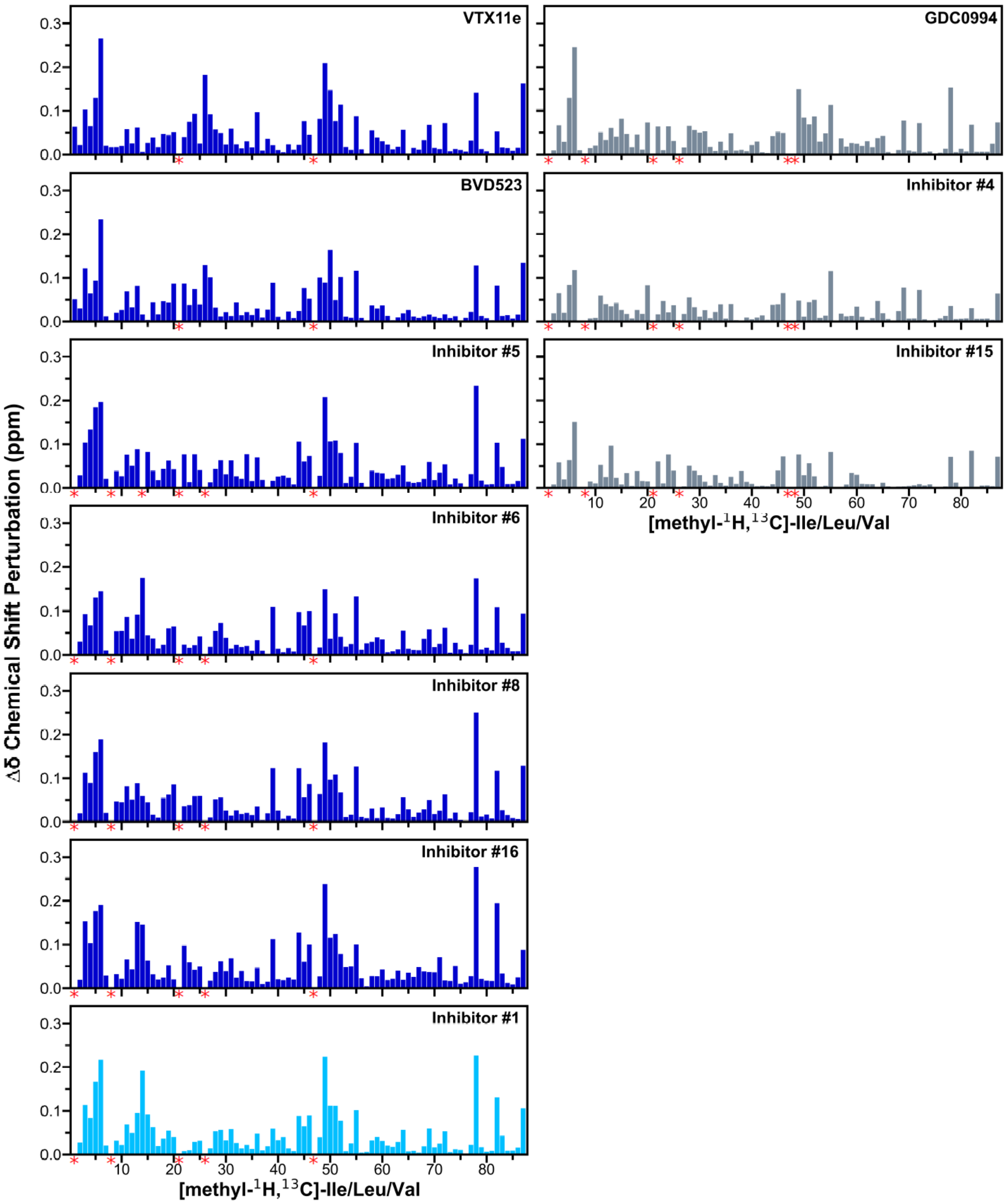


**Figure S10. Chemical shift changes induced by inhibitor binding to 2P-ERK2.** 2D-HMQC spectra collected on [methyl-$^{13}$C,$^{1}$H]-ILV labeled 2P-ERK2 complexed with VTX11e, BVD523, GDC0994, and the selected subset of seven inhibitors in **Fig. 6**. Chemical shift perturbations (CSP) were calculated by the equation: $\Delta\delta$ (ppm) = SQRT[$(\delta_H^2 + 0.20(\delta_C^2))$], and plotted for each ILV residue methyl. Red asterisks below the x-axes indicate methyl peaks that could not be traced in the ligand-bound state, ascribed to peak broadening. Numberings of methyl assignments, their chemical shifts and CSP calculations are listed in **Suppl. Dataset S2**.

### Figure S11

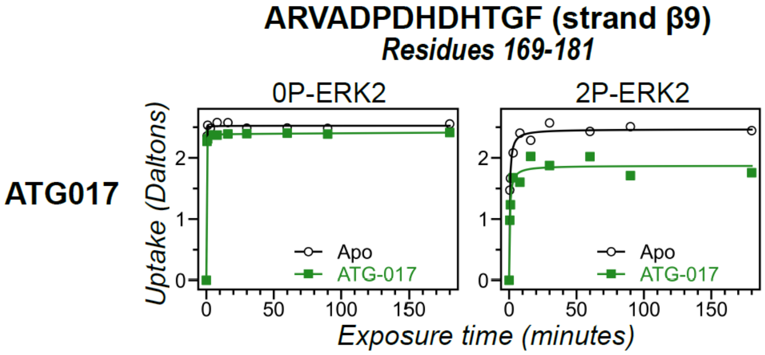


**Figure S11. HDX time courses responsive to ATG017 binding.** HDX time courses for strand β9 (peptide 169-181, ARVADPDHDHTGF). Open symbols show deuterium uptake in 0P- or 2P-ERK2 apoenzymes. Closed symbols show deuterium uptake in 0P- or 2P-ERK2 complexed with ATG017 (green). ATG017 decreases HDX uptake into strand β9 to a larger degree compared to GDC0994, similar to the behavior of the DFG motif, P+1 and helix αF (**Fig. 9D**).

### Figure S12

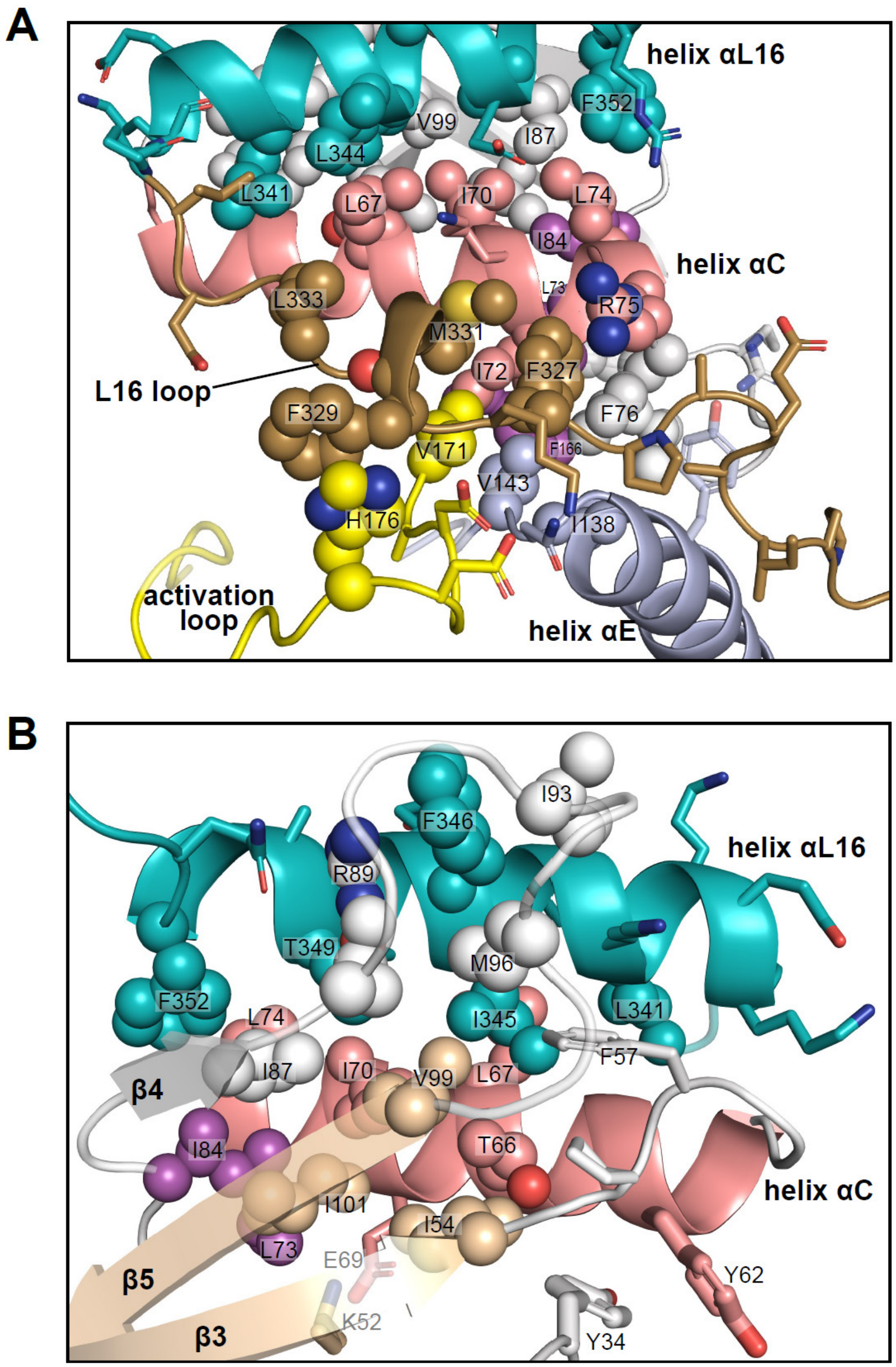


**Figure S12. Interactions of L16 and helix αL16 with N-lobe elements.** Views of 2P-ERK2 (PDBID:2ERK), showing hydrophobic residue interactions between **(A)** loop L16 (M331, F327) and helices αC and αE (I72, R75, F76, I138). Residues in helix αC in turn form second sphere interactions with residues nearby catalytic residues in β3 (I54, nearby K52), β8-β9 (F166, DFG motif), and β6 (V143, nearby HRD). **(B)** Residue interactions showing close connections between helix αL16 (I345, F346, T349, F352), and helix αC (I54, F57, T66, L67, I70, L74, I84, I87) and loop β4-β5 (R89, I93, M96, V99). Residues in magenta (L73, I84, F166) participate in the R-spine in ERK2.

#### Figure S13

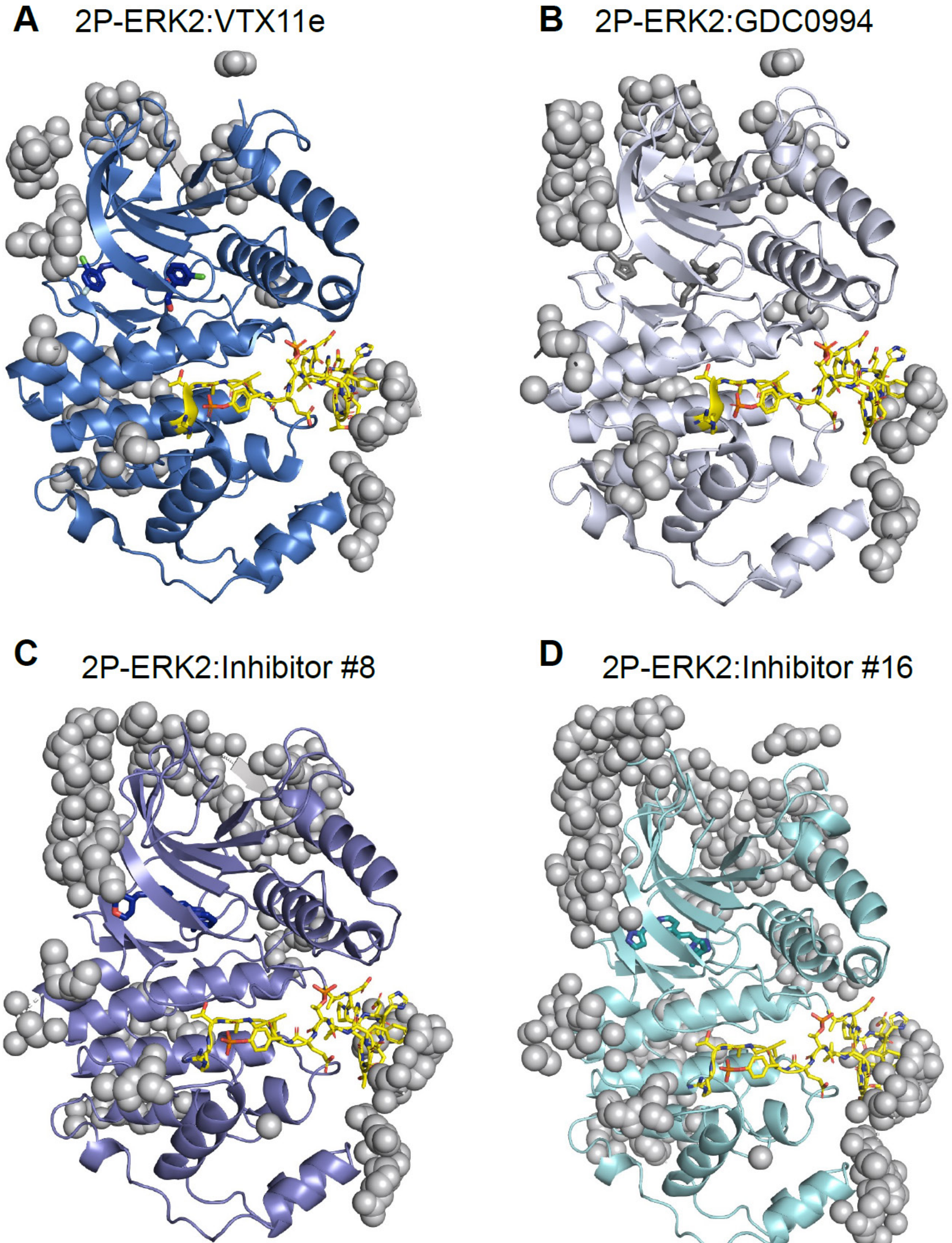


**Figure S13. Crystal contacts with the activation loop in X-ray structures of ERK2.** Structures of **(A)** 2P-ERK2:VTX11e (PDBID:6OPK), **(B)** 2P-ERK2:GDC0994 (PDBID:6OPH), **(C)** 2P-ERK2:Inhibitor#8 (PDBID:8U8K), and **(D)** 2P-ERK2:Inhibitor#16 (PDBID:8U8J). The activation loop is rendered in yellow, and atoms to the asymmetric unit from protein neighbors within 5 Å are shown as grey spheres.
